# Lineage recording of zebrafish embryogenesis reveals historical and ongoing lineage commitments

**DOI:** 10.1101/2020.07.15.203760

**Authors:** Zhuoxin Chen, Chang Ye, Zhan Liu, Shanjun Deng, Xionglei He, Jin Xu

## Abstract

It has been challenging to characterize the lineage relationships among cells in vertebrates, which comprise a great number of cells. Fortunately, recent progress has been made by combining the CRISPR barcoding system with single-cell sequencing technologies to provide an unprecedented opportunity to track lineage at single-cell resolution. However, due to errors and/or dropouts introduced by amplification and sequencing, reconstruction of accurate lineage relationships in complex organisms remains a challenge. Thus, improvements in both experimental design and computational analysis are necessary for lineage inference. In this study, we employed single-cell Lineage tracing On Endogenous Scarring Sites (scLOESS), a lineage recording strategy based on the CRISPR-Cas9 system, to trace cell fate commitments for zebrafish larvae. With rigorous quality control, we demonstrated that lineage commitments of complex organisms could be inferred from a limited number of barcoding sites. Together with cell-type characterization, our method could homogenously recover lineage information. In combination with the cell-type and lineage information, we depicted the development histories for germ layers as well as cell types. Furthermore, when combined with trajectory analysis, our methods could capture and resolve the ongoing lineage commitment events to gain further biological insights into later development and differentiation in complex organisms.

## Introduction

Resolving the lineage relationships among cells from a zygote to understand the process of cell proliferation and differentiation is of great interest in the field of developmental biology. So far, only the lineage of nematode *Caenorhabditis elegans* has been successfully tracked at single-cell resolution (***Deppe et al., 1978***; ***Sulston et al., 1983***; ***Sulston and Horvitz, 1977***). There has been little progress in non-eutelic, multicellular organisms due to their large number of cells and opacity. Recently, by combining single-cell RNA-sequencing (scRNA-seq) with the CRISPR-Cas9 system, many researchers have developed new strategies to simultaneously record the lineage histories and characterize cell types. These approaches have provided unprecedented opportunities to understand the development and physiology of different organisms (***Alemany et al., 2018***; ***Chan et al., 2019***; ***Raj et al., 2018***; ***Spanjaard et al., 2018***).

One challenge in utilizing this system is the number of scars created by the CRISPR system is limited compared to the huge number of cells in an individual. In addition, technical issues, such as low recovery rates of barcoding sites, recurring mutations and chimeric reads, are barriers for accurate lineage relationship reconstruction (***Spanjaard et al., 2018***; ***Wang and Wang, 1997***). Further optimization of experimental design and data processing is essential to improve the signal-to-noise ratio.

In our previous study, we proposed a method using the CRISPR-Cas9 system to edit endogenous sites for lineage recording, which has higher recording capacity and recovery rates than previous methods (***Ye et al., 2020***). In this study, we applied this method (called scLOESS) to track the lineage of cells from 7dpf zebrafish larvae. With rigorous quantification, we demonstrated that the cell lineage commitments of complex organisms can be inferred from a limited number of barcoding sites.

## Results

### Characterization of cell types and assignment of germ layers

We injected Cas9 mRNA and the 78-sgRNA pool into zebrafish embryos as previously described (***Ye et al., 2020***). At 7 days post-fertilization (dpf), an injected embryo was dissociated into singlecell suspension and subjected to scRNA-seq as well as barcoding-site amplification (***Figure 1A***). We obtained more than 12,000 cells with an average of approximately 3,000 unique molecular identifiers (UMIs) and 900 genes per cell. Using an unsupervised clustering approach, we identified 41 clusters and assigned them to particular cell types based on their feature genes (***Figure 1B-F*** and ***Figure 1–source data 3***) (***Butler et al., 2018***). Based on information from the Zebrafish Information Network (ZFIN) database and previous studies, we further classified these clusters into four different germ layers (***Howe et al., 2013***; ***Thisse and Thisse, 2008***). Notably, we identified almost all the cell types in zebrafish larvae, suggesting that targeting endogenous sites does not noticeably affect normal development.

**Figure 1.**
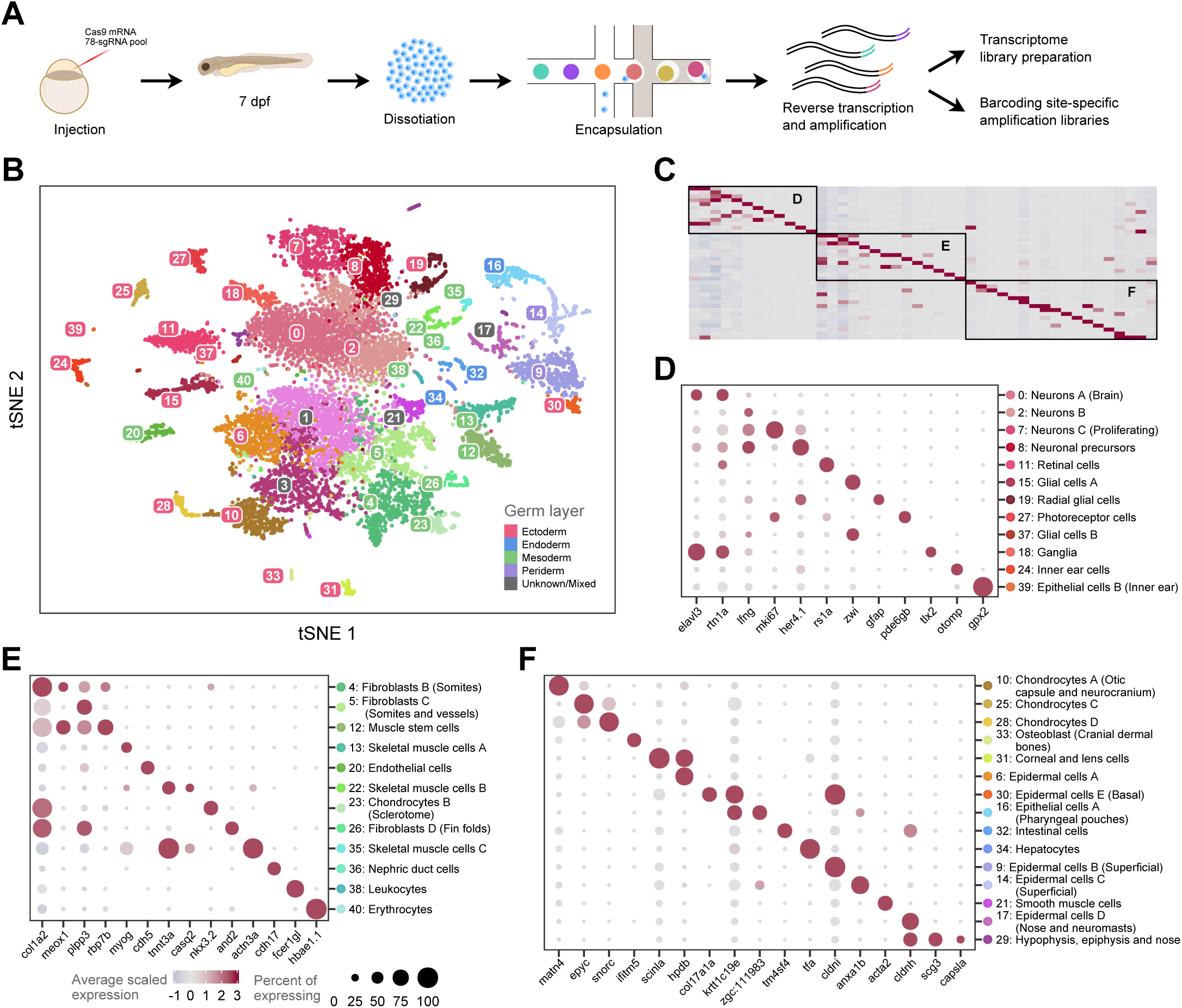
Cell-type assignment by scRNA-seq. (**A**) Schematic of simultaneous detection of single-cell transcriptome and lineage information. (B) t-SNE plot of 12,975 cells from zebrafish larva (7 dpf) clustered into 41 groups. Based on the differentially expressed genes, each group was assigned to different cell types and germ layers, as indicated by the color code on the y-axis (D to E) and the legend. Cell types with unknown or mixed origins are labeled in gray. (**C**) Heatmap of scaled expression of representative marker genes for each cell type. The detailed information is shown in (**D**-**E**). (**D**-**E**) Dot plot of representative marker genes of diverse cell types from the neural tube (including posterior placodal area), mesoderm, and others, respectively. Dot size denotes the fraction of cells expressing the marker genes and the color indicates the average scaled expression level. **Figure 1–Figure supplement 1.** The distribution of UMI number per cell with different marker genes. **Figure 1–source data 1.** Targets. **Figure 1–source data 2.** Primers. **Figure 1–source data 3.** Maker genes of various cell types.

### Lineage barcode extraction and tree reconstruction

With cell-type characterization, we investigated the mutations on the barcoding sites to reconstruct lineage relationships among cells. To obtain accurate lineage information, we selected the barcoding sites detected in more than 80% of the cells and mutations that overlapped with the 6 base pairs upstream of the PAM sequences (***Figure 1–source data 1***). We further filtered potential sequencing errors, such as chimeric reads and doublets as well as recurring mutations, to reduce the technical noise (see Materials and Methods). After obtaining high-quality mutations, we investigated the power of our recording system. The confident mutations created on the barcoding sites were referred to as scars. We found that there were 512 distinct scars in total with various lengths of insertions and deletions, which is consistent with the mutation spectrum edited by Cas9 (***Figure 2A***). We found that scars covered 70% of the cells without cell-type and germ-layer bias, and more than one scar was detected in more than half of the cells (***Figure 2B-C***). These results revealed that our method homogenously marked cells after quality control, which is essential for a highly informative and accurate lineage recording system.

**Figure 2.**
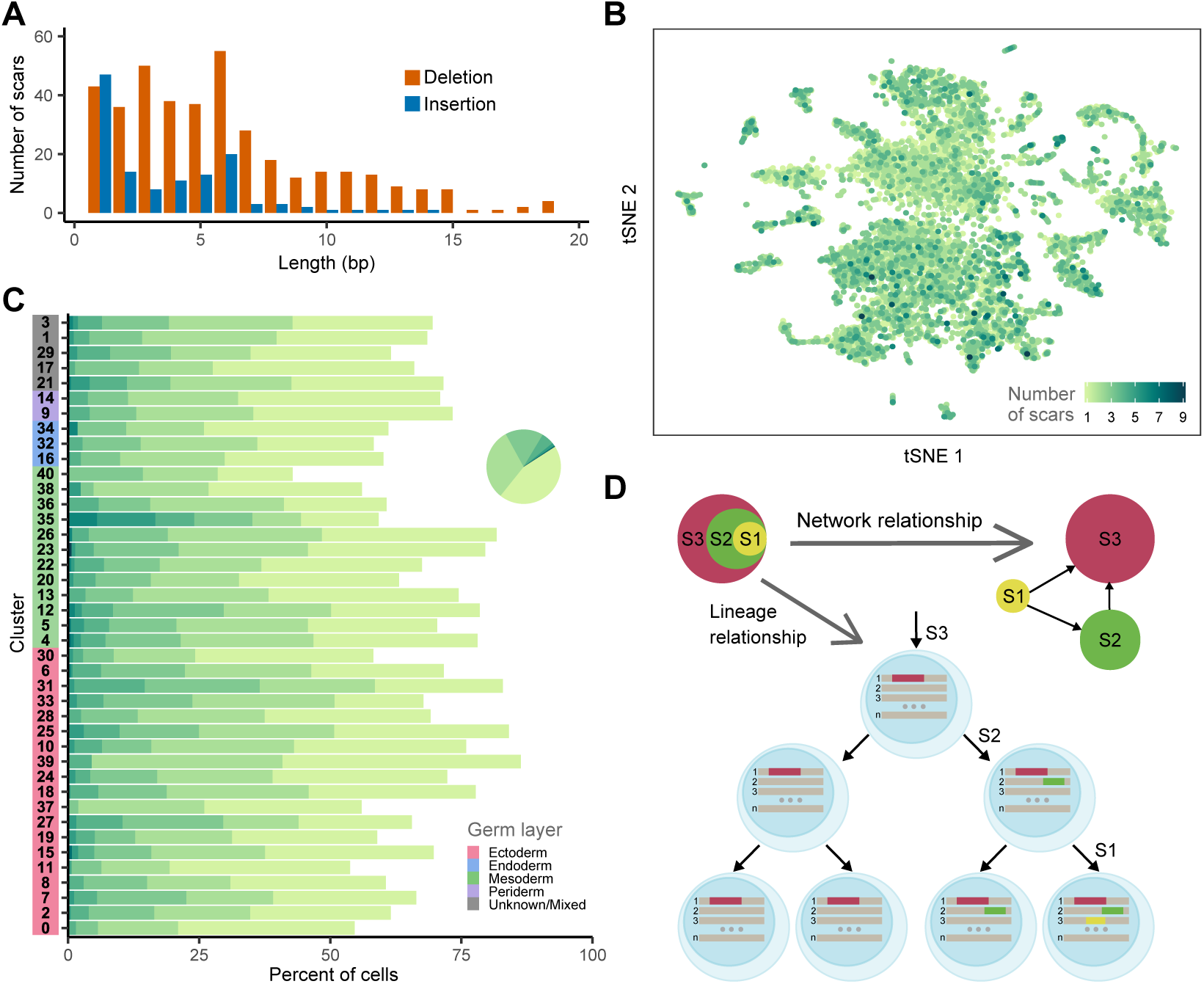
Lineage barcoding in zebrafish larvae by scLOESS. (**A**) Length distribution for deletions and insertions of various scars. (**B**) t-SNE representation of cells colored by the number of detected scars. Most of the cells were marked by more than one scar, indicating the ability of broadly endogenous barcoding of scLOESS. (**C**) Proportion of cells with a different number of scars within each cell type. The colors in the y-axis indicate the germ layer from which the cells originated. Color code for the number of scars is as indicated in (B). Pie chart shows the fractions of cells with a different number of scars. (**D**) Example of lineage tree reconstruction from unambiguously placed scars. For all pairs of scars, their inclusion relationships can be illustrated using a network graph and then reconstructing the lineage relationships among cells based on their scar profiles. **Figure 2–Figure supplement 1.** Network relationships of non-recurring scars. **Figure 2–Figure supplement 2.** The reconstructed lineage tree for 7 days post-fertilization zebrafish larva. **Figure 2–Figure supplement 3.** Proportions of less frequent scars per site in individual cells with various number of scars. **Figure 2–Figure supplement 4.** Scar networks before filtering.

We next reconstructed the lineage relationships using scar network graphs (***Spanjaard et al., 2018***). Basically, the scars that were created in a descendant cell marked by another scar could be detected simultaneously with the latter, and the historical relationships of scars could be inferred from the network (***Figure 2D***). We reconstructed the lineage tree according to the final network graph (***Figure 2–Figure Supplement 1***) and assigned cells to the terminal nodes in the tree based on their scar profiles (see Materials and Methods). The reconstructed lineage tree revealed that the cell fates of progenitors become restricted to certain cell types in the same germ layer during embryogenesis (23 out of 85 terminal nodes, ***Figure 2–Figure Supplement 2***). The reconstructed tree comprised only approximately 10% of the barcoded cells, which was not informative enough for us to have a closer examination at the lineage relationships between diverse cell types. Although we could not accurately order all the scars in a lineage tree, cells with identical sets of scars were very likely to have been developed from the same recent ancestor.

### Reconstructed lineage relationships reveal developmental histories

We next investigated whether the lineage information recapitulated more comprehensive developmental histories in zebrafish. To depict accurate lineage relationships among different cell types, we further excluded scarring paths that may have been created right before differentiation (see Materials and Methods). As expected, most of the remaining scarring paths (>75%) were germ-layer-restricted. Compared to random sampling based on the number of cells in each germ layer and scarring path (***Figure 3–Figure Supplement 1A***), the observed number of path-marked cells within a single germ layer was substantially larger than expected (Figure 3A). We also noticed that in germ-layer-specific paths, most cell types were connected to multipotent stem cells (cluster 4 for mesoderm is shown in ***Figure 3B***, cluster 7 for ectoderm is shown in ***Figure 3–Figure Supplement 1B***). These results indicate that during development, progenitors give rise to both differentiated cells and stem cells that retain their capacity to differentiate into other cell types. To further quantify the lineage connection between cell types, we counted and normalized the connections between each pair of cell types in the filtered scarring paths and presented the lineage relationships using a circular plot (***Figure 3C***). We found that cells originating from the same regions displayed a higher degree of common ancestry, which agrees with the developmental history (***Kimmel et al., 1990***). These results demonstrate the power of our method to recapitulate the cell fate commitments during early development in zebrafish.

**Figure 3.**
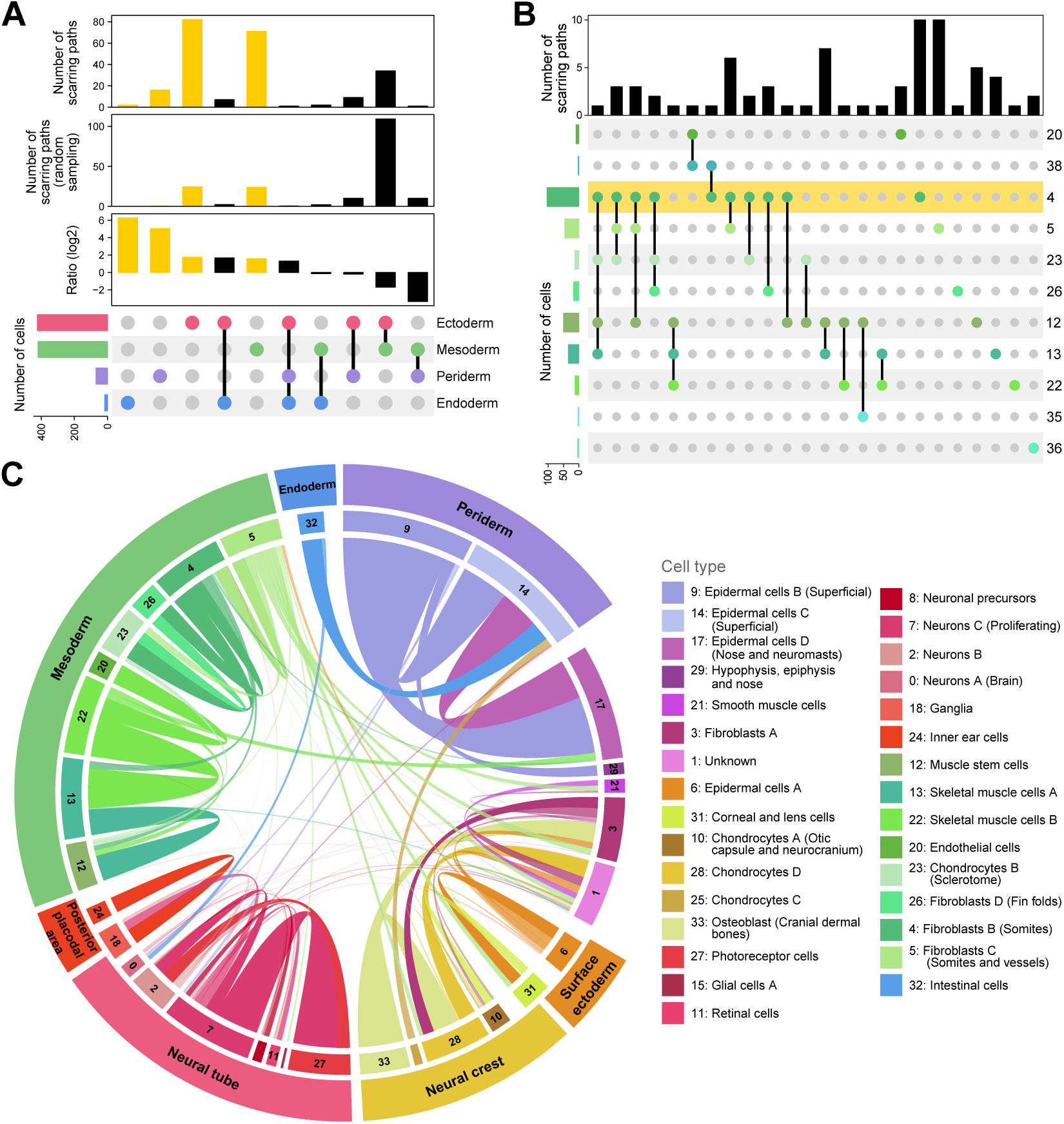
Lineage relationships between different cell types. (**A**) Upset plot showing the number of scarring paths detected within or between distinct germ layers. The top two bar plots showing the number of scarring paths from our data (observed) and random sampling, based on the cell number of each germ layer (expected). The bottom bar plot showing the ratios (log2 scale) between the observed and expected number of paths. Scarring paths containing cells from unknown/mixed germ layers are removed. Yellow bars indicate scarring paths that were only detected in cells from one germ layer. The color code for the germ layers is as annotated in ***Figure 1B***. (**B**) Upset plot for the number of scarring paths detected within or between cell types from the mesoderm. The color code for cell types is as annotated in ***Figure 1E***. The yellow shade represents multipotent stem cells. (**C**) Chord diagram for relative lineage connection strengths, in which links between cell types represent lineage connections. The link widths are proportional to the ratio of the observed connections to all possible connections between each pair of cell types. Connections between cells within the same cell type are removed for clarity. The alpha transparencies of links are scaled by their ranks (1 for the top 25%, 0.75 for the top 25−50%, 0.5 for the top 50−75%, and 0.25 for the rest). The ectoderm is further divided into the neural tube, neural crest, posterior placodal area, and surface ectoderm. **Figure 3–Figure supplement 1.** Lineage information for simulation. **Figure 3–Figure supplement 2.** Fin muscle cells share a close relationship with endothelial cells. **Figure 3–Figure supplement 3.** Comparison of various methods for single-cell lineage tracing in vertebrates. **Figure 3–Figure supplement 4.** The relationship of the number of cell types and the proportions of the top two cell types.

We found that the most prominent connection was in periderm (also known as the enveloping layer (EVL); cluster 9, 14), which is the earliest cell commitment event in zebrafish development (***Haddon and Lewis, 1996***; ***Kimmel et al., 1990, 1995***; ***Lee et al., 2014***; ***Teixeira Rosa et al., 2019***). We also noticed that epidermal cells of the nose (cluster 17) had a strong lineage relationship with cells from the periderm, suggesting that a portion of those cells were descendants of the periderm. Nevertheless, there was an unexpected connection between intestinal cells (cluster 32) and epidermal cells C (cluster 14), which was due to incorrect clustering and annotation of intestinal cells, according to the selected feature genes (***Ivanova et al., 2015***).

Furthermore, the connections between different cell types provided detailed information on lineage commitment during development. For example, a portion of skeletal muscle cells (cluster 22) and endothelial cells (cluster 20) shared a common developmental origin (***Figure 3C*** and ***Figure 3–Figure Supplement 3***). Inspection of the expression signature of these cells revealed that they are posterior cardinal vein endothelial cells (*lyve1a*^+^) and pectoral fin muscle cells (*casq2*^+^), both of which are derived from the posterior lateral plate mesoderm (PLPM) (***Cleaver and Krieg, 1998***; ***Fouquet et al., 1997***; ***Hamburger and Hamilton, 1951***; ***Martin, 1998***; ***Zhong et al., 2001***). Moreover, there has been controversy as to whether the trunk neural crest contributes to the fin mesenchyme (***Green et al., 2015***). Our results showed that fin fibroblasts (cluster 26) shared a common lineage origin with cells from the paraxial mesoderm (cluster 4, 23) instead of the neural crest, which was consistent with a recent study by ***Lee et al.*** (***2013***).

The neural crest is a migratory embryonic cell population that gives rise to a wide variety of cell types, including the craniofacial skeletal cells, cornea cells, and smooth muscle cells (***Akula et al., 2019***; ***Etchevers et al., 2001***; ***His, 1868***; ***Kague et al., 2012***). Remarkably, our results showed that cluster 31, comprising cornea and lens cells, shared close lineage relationships with cells from the neural crest (cluster 10) and surface ectoderm (cluster 6), which was in agreement with the organogenesis of vertebrate eye (***Greiling and Clark, 2009***; ***Langenberg et al., 2008***; ***Soules and Link, 2005***; ***Tamm, 2011***; ***Yoshikawa et al., 2007***). Our results also revealed that cells derived from the posterior placodal area, another thickening of the ectoderm, were more closely related (***Ladher et al., 2010***). As shown in ***Figure 3C***, ganglia (cluster 18), which contain the statoacoustic (VIII) ganglia and epibranchial ganglia, had a shared lineage origin with inner ear cells (cluster 24). This finding was also consistent with previous studies (***Freter et al., 2008***; ***Haddon and Lewis, 1996***; ***Sun et al., 2007***; ***Wikstrom and Anniko, 1987***).

Collectively, these results demonstrate that our method can decipher the developmental histories of thousands of cells during embryogenesis.

### Tracking ongoing lineage commitment

As cell proliferation and differentiation are still ongoing at 7 dpf, we wondered whether we could detect current lineage commitment events using our strategy. During ontogenesis, progenitor cells become biased toward certain fates by changing their transcriptional states. It is known that mesenchymal cell (fibroblast) fate commitment in the paraxial mesoderm is still ongoing during the larva stage (***Hollway et al., 2007***; ***Lleras Forero et al., 2018***). Therefore, we applied pseudo-time analysis based on expression signatures to predict the developmental trajectories. We first analyzed the cell trajectories of fibroblasts in the paraxial mesoderm and then investigated how these cell trajectories supported by their lineage information (***Figure 4A***) (***Crotwell and Mabee, 2007***; ***Maradonna et al., 2013***; ***Morin-Kensicki and Eisen, 1997***). We found that the progenitor cells in the paraxial mesoderm would gradually acquire fates of either the sclerotome (*nkx3.2*^+^, *matn3b*^+^), myotome boundaries (*COMP*^+^), or fin fold mesenchyme (*and2*^+^) (***Figure 4B-D, Figure 4–Figure Supplement 1***) (***Crotwell and Mabee, 2007***; ***Ko et al., 2005***; ***Zhang et al., 2010***). Cells with the same scarring path matched either single (4 in 16 scarring paths with more than 5 cells) or multiple trajectories (12 in 16 scarring paths) (***Figure 4B-D***).

**Figure 4.**
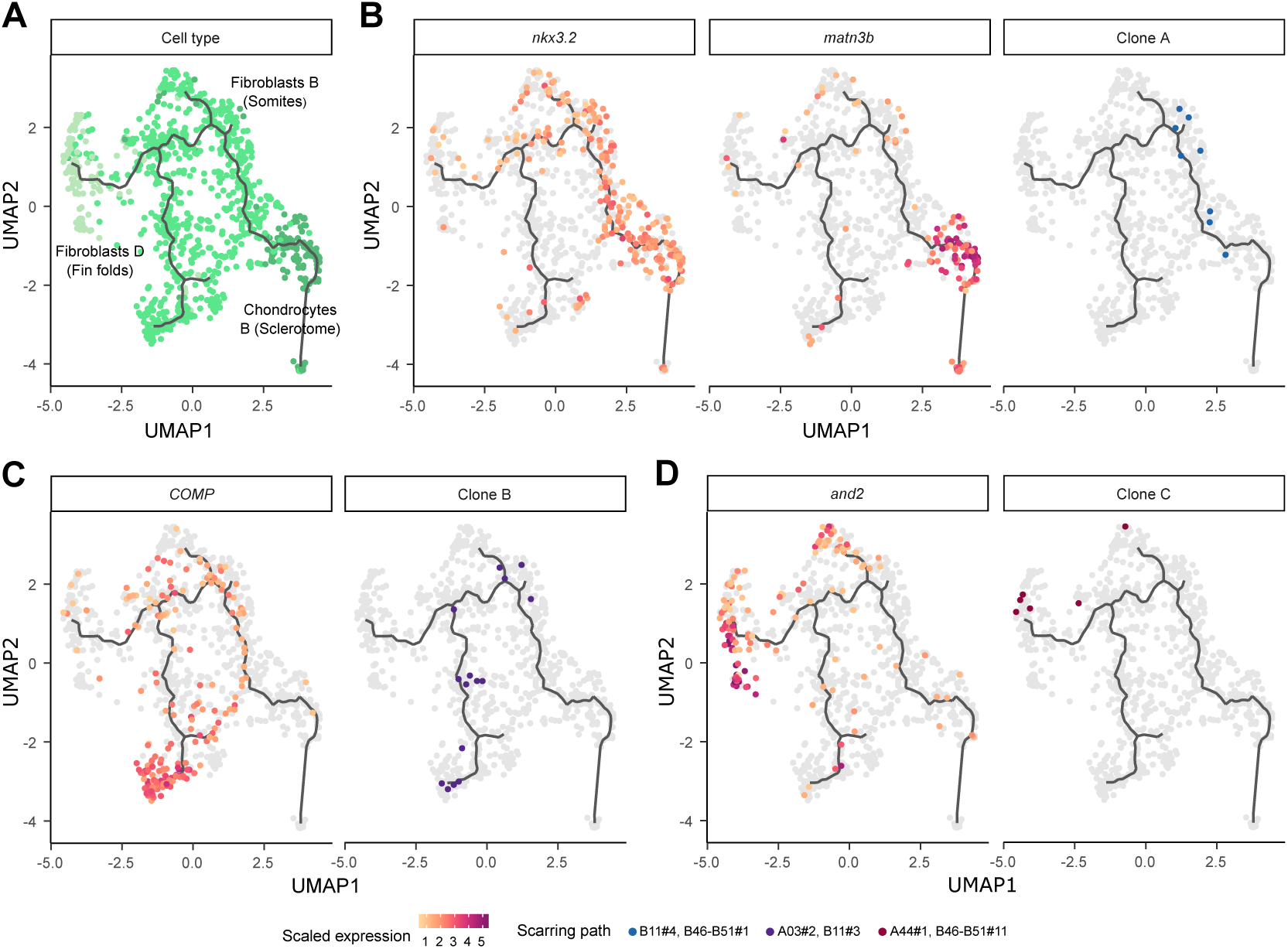
Scarring paths track ongoing lineage commitment. (**A**) UMAP visualization of cells in the paraxial mesoderm with the inferred trajectories generated by Monocle3. Color code for the cell types is as in ***Figure 1***. (**B-D**) Expression patterns of marker genes for the sclerotome (*nkx3.2* and *matn3b*), myotome boundaries (*COMP*), or fin fold mesenchyme (*and2*), as well as the cells marked by the same scarring path along the trajectories. **Figure 4–Figure supplement 1.** Transcriptional transition along different trajectories. **Figure 4–Figure supplement 2.** Site frequency spectrum of different trajectories.

However, the number of cells with the same scarring path is too limited to tell how these progenitor cells determine their fates, when there are multiple routes. We further estimated the effective population size of each trajectory, based on the principle from population genetics (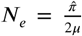 for haploidy genome). The genetic diversity 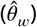 of each cell population was estimated from the scarring paths under the infinite sites model (***Kimura, 1971, 1969***). Interestingly, the genetic diversity of cells within the trajectory toward fin was the lowest (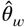 for the myotome, sclerotome, and fin were 7.69, 6.41, and 4.76, respectively) (***Watterson, 1975***). As the scaring path was created earlier, we assumed the mutation rate is equal for each population and deduced that the population size of the progenitors contributed to fin fold mesenchyme was 62% of the myotome (***Figure 4–Figure Supplement 2***). It has been reported that fin fold mesenchyme mainly derives from the marginal region of the paraxial mesoderm, which may reduce the effective progenitor population size (***Lee et al., 2013***). Taken together, our results indicate fibroblasts may commit to different fates stochastically, while spatial and signaling information can shape their behavior and lineage potential.

In summary, our results confirmed the ability of scLOESS to capture and resolve the ongoing lineage commitment events and gain further biological insights into the development and differentiation of complex organisms.

## Discussion

The application of scLOESS in zebrafish larvae revealed that it was powerful enough to decipher cell fate commitment during the early development of complex multicellular organisms. Taking advantage of distinguishable endogenous barcoding sites, our method exhibited higher recording capacity and recovery rates than previous methods (***Figure 3–Figure Supplement 3***). We demonstrated that accurate lineage relationships could be reconstructed from dozens of high-quality barcoding sites, which was consistent with the simulations in a previous study (***Salvador-Martínez et al., 2019***). This was achieved by filtering out less homogeneously captured barcoding sites and potential chimeric reads (see Materials and Methods), thus improving the performance of our strategy. Future incorporation of more barcoding sites could facilitate the reconstruction of a higher-resolution fate map.

As mentioned above, scLOESS not only recapitulated cell ancestry, but also shed light on temporal relationships of cell differentiation. Although our method exhibited several advantages, limitations were also notable. Reconstruction of accurate lineage trees was hampered because of the stochasticity of mutation generation by the CRISPR system, which was uncoupled with the cell cycle and resulted in recurring scar formation. Moreover, lack of a continuous barcoding system in our design hindered investigation of later developmental stages. Hence, future incorporation of scLOESS into an inducible system may be necessary to unravel relationships at various timepoints.

Although targeting endogenous sites potentially could have affected the normal development of zebrafish, we identified most of the cell types in our scRNA-seq data, implying that disruption of normal development was not a major concern. However, it is worth noting that the number of lens cells was relatively small compared with previous studies (***Spanjaard et al., 2018***). With a careful examination of the raw data, we found that this was due to their overall low expression levels (***Figure 1–Figure Supplement 1***), rendering them unable to pass the initial threshold (>500 UMIs) for downstream analysis. Nevertheless, we did not attempt to retrieve all the lens cells because they most likely had few reads on our barcoding sites. This suggests that our method may not be suitable for tracking lineages with low expression levels.

Further combination of our method with spatial scRNA-seq technology may help to illustrate more comprehensive and accurate fate mapping because clusters uncovered by droplet-based scRNA-seq comprise cells from mixed origins (***Figure 1*** and ***Figure 3C***). Thus, we anticipate that scLOESS will lead us toward fine-resolution, spatiotemporal fate mapping of vertebrate development.

## Methods and Materials

### Zebrafish microinjection and library preparation

Wild-type (AB) zebrafish were maintained at 28.5°C under a 14-h light/10-h dark cycle. The Cas9 mRNA and 78-sgRNA pool were generated by in vitro transcription, as described previously (***Ye et al., 2020***). For high-information lineage tracing, we injected one-cell stage embryos with a mixture of 2 nl Cas9 mRNA (final concentration 350 ng/µl) and the 78-sgRNA pool (final concentrations 585 ng/µl and 7.5 ng/µl, respectively).

Single-cell suspension was prepared from a 7 dpf zebrafish larva, which was filtered through a 35-µm strainer and subjected to the 10× Genomics Chromium Single Cell 3’ Library & Gel Bead Kit v2, with minor modifications to the manufacturer’s instructions. Specifically, after the first cDNA amplification (a total of 9 amplification cycles) and cleanup procedures, the amplification product was divided into two equal aliquots. One aliquot was fragmented and used for the conventional scRNA-seq library preparation. The other aliquot was reserved for target-specific amplification experiments, consisting of three replicates. Explicitly, 1 µg of cDNA library was first amplified for 15 cycles using the cDNA_READ1 and cDNA_TSO primers (***Figure 1–source data 2***). Afterwards, the amplicons were purified and split into 90 equal aliquots for site-specific amplification, in which each site was amplified for another 25 cycles with the cDNA_READ1 and cDNA_target primers. Then, the PCR products were mixed in equimolar amounts after purification, followed by the addition of Illumina compatible adaptors for NGS sequencing. All PCR reactions were performed using Q5 High-Fidelity DNA Polymerase, and the primer sequences used in this study are listed in ***Figure 1–source data 2***. Next, both the scRNA-seq and the amplicon library were analyzed using the Agilent 2100 Bioanalyzer and sequenced on the Illumina HiSeq X platform (PE150 mode), while incorporating 30% of the spike-in DNA.

All animal experiments were performed in accordance with the Animal Research and Ethics Committees of Sun Yat-Sen University and the National Guidelines for the Care and Use of Laboratory Animals (China).

### Cell type identification

The resulting transcriptome library was mapped to the reference genome (GRCz11) using Cell Ranger 3.1.0, and cells with fewer than 500 UMIs were excluded. Then, we used the Seurat version 3.1.2 for clustering and identification of differentially expressed genes as described below (***Butler et al., 2018***). We first discarded genes that were found in fewer than three cells as well as cells that expressed fewer than 200 of those genes. Next, gene expression was log-normalized and the top 2,000 most variable genes were selected for principal component analysis. Using the Louvain algorithm, cells were clustered into 41 groups with the top 40 components and a resolution of 1.8. Finally, the differentially expressed genes were identified for each cluster (FindAllMarkers function, only.pos = TRUE, min.pct = 0.25, logfc.threshold = 0.3), and cell-type information was assigned to each cluster based on the top 50 differentially expressed genes (***Figure 1–source data 3***), based on the knowledge from literature and the ZFIN database (***Howe et al., 2013***). Information regarding the germ layers from which different cell types were derived was also carefully retrieved.

### Scar analysis

Together with the three barcoding site-specific amplification libraries and the standard transcriptome library, we had four replicates for scar identification. The site-specific libraries were also mapped to the reference genome using Cell Ranger 3.1.0. We first filtered out reads with uncorrected cell barcodes that were not in the filtered cell barcode matrix. Then, reads that could be mapped to the barcoding sites were extracted for each site, respectively, and mutated reads (with CIGAR strings containing “I/D”) were deduplicated and merged using CAP3 and then mapped to the reference target sequences using MAFFT version 7.402 (***Huang and Madan, 1999***; ***Katoh and Standley, 2013***). The reference sequences were manually trimmed to the shortest blocks that covered all the indels that overlapped with the barcoding sites. This step mitigated the impact of sequencing errors surrounding the barcoding sites in downstream analysis. Afterwards, the information for replicate identification, cell barcodes, and UMI sequences of each sequence were extracted. Then, the reads were mapped to the reference sequences using T-COFFEE (Version_11.00) for more accurate alignment and were transformed into a four-letter sequence, as described previously (***Perli et al., 2016***). Aligned sequences were only considered to the PAM sequence for each site. Because heterogeneously presented barcoding sites (sequencing dropouts) can influence the accuracy of lineage reconstruction, we excluded sites that were detected in less than 80% of the cells; therefore, 49 barcoding sites remained.

For all the droplet-based single-cell sequencing methods, there are three major errors that can affect the accuracy of scar analysis: amplification and sequencing errors, chimeric reads, and doublets/multiplets (***Wang and Wang, 1997***). Thus, we performed multiple filtering steps to mitigate the impact of these errors, according to the following criteria. 1) If there were wild-type and mutant sequences for the same UMI in the same cell, we selected the sequences with the highest replicate counts. 2) If there was more than one scar for the same UMI, then the scar with the most replicates in that cell or with the fewest mismatches was chosen for downstream analysis. 3) For each cell, if there were scars with the same core scar (23 bp) but different full scars (trimmed block) of the same length, then they were considered as the same. 4) We further merged scars that had Hamming distances of 1, unless there was a single-nucleotide polymorphism (SNP) surrounding the barcoding site (see ***Figure 1–source data 1***). 5) We incorporated scars that were only detected in one cell into scars that were detected in multiple cells that had only one mismatch.

After completion of the above-mentioned filtering steps for sequencing errors, we counted the UMI numbers for each scar in individual cells. Although the proportions of chimeric reads could not be inferred from the wild-type alleles, we estimated it from the cells that had two or more scars. If there was more than one scar in a cell, it most likely resulted from chimeric reads and/or doublets/multiplets because the possibility of creating mutations in both alleles was low. Thus, we calculated the proportions of less frequent scars per site in individual cells (***Figure 2–Figure Supplement 3***). Most of the less frequent scars (75% quantile) accounted for less than 9% of the total number of UMIs per site; therefore, we filtered out scars that had less than 9% of the total number of UMIs for each site in single cells. Cells that still had at least two scars in the same barcoding site (probably doublets/multiplets) as well as scars that were detected in fewer than three cells were also excluded. Finally, we ranked the scars according to their frequencies in the barcoding sites, which was used as their identification (e.g., A03#2).

### Tree reconstruction

To reconstruct an accurate lineage tree, we first calculated the connections of scars. For all pairwise combinations of scars that were observed in this study, scar S1 was considered to be created next to scar S2 if the number of cells marked by it was smaller than that of scar S2 and most of the cells marked by scar S1 were also marked by scar S2, after considering potential dropouts. Thus, we calculated the percentage of overlaps between scars S1 and S2, and if the overlap was greater than 60% and supported by at least three cells, then it was considered that there was a connection between them. This was because the average recovery rate of filtered barcoding sites was 96%. Therefore, if the recovery rates of each allele were independent, then the possibility of detecting each barcoding allele could be calculated by 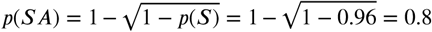, where *p*(***S***) was the possibility of detecting the barcoding site. Thus, if two scars had a connection (meaning one scar was created after another), the chance of detecting both scars in the descendant cell was approximately 60%.

Next, because the CRSIRP-Cas9 system may generate recurring mutations, we filtered out scars that could have been created multiple times. As shown in ***Figure 2–Figure Supplement 4A***, if one scar belonged to two scars and there was no connection between them, then it was a recurring scar. The initial scar network is shown in ***Figure 2–Figure Supplement 4B***. After removing potential recurring scars, there were 42 sub-networks left, representing 42 branches. Thus, according to the final scar network (***Figure 2–Figure Supplement 1***), we reconstructed the lineage tree of the zebrafish larva and cells were placed within the terminal nodes based on their scar profiles.

### Lineage relationship reconstruction

We attempted to test if our method could unravel the canonical lineage relationships among different cell types. We took all the barcoded cells into account and grouped them based on their scar profiles. Distinct scar profiles were considered to stand for unique lineage clades, and cells with identical scar profiles were very likely developed from the same recent ancestor. Therefore, we referred to the scar profiles as scarring paths, and a lineage connection was considered to exist between a pair of cell types if they were in the same scarring path. First, we selected scarring paths that contained more than one cell and that were significantly biased toward particular cell types (one-sided Fisher’s exact test, FDR < 0.05). The progenitors marked by the filtered paths were expected to restrict their developmental potential to certain cell types. Next, to further constrain the timepoints marked by the scarring paths, we discarded the scarring paths with the top two cell types that accounted for less than 70% of the cells within them. The rational for this was that if a scarring path was created right before differentiation, then progenitors marked by that scarring path should give rise to a limited number of cell types. As shown in ***Figure 3–Figure Supplement 4***, if the proportion of the top two cell types in a path was less than 70%, then the path was likely marked by cell types from multiple germ layers.

Thus, cell types in the same filtered path were expected to share a closer lineage relationship. To reconstruct accurate lineage relationships, we summed up the connections between each pair of cell types in the filtered paths and discarded connections supported by fewer than three cells of either cell type. Finally, the connection strength was calculated using the ratio of observed connections to all possible connections between the cells of each cell-type pair. More precisely, the total connection number observed in our data between cell types A and B was calculated by 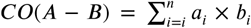 where *a*_*i*_ and *b*_*i*_ mean the number of cells from cell type A and B in path *i*, respectively. Furthermore, the number of all possible connections between cell type A and B was calculated by 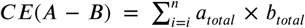, where *a*_*total*_ and *b*_*total*_ mean the total cell number in each cell type. Thus, the connection strength of cell type A and B was calculated by ***P*** (***A*** − ***B***) = ***CO***(***A*** − ***B***)/ ***CE***(***A*** − ***B***).

### Differentiation trajectory analysis

With the top 100 components and a resolution of 0.01, cells from the paraxial mesoderm (cluster 4, 23, 26) were clustered and ordered in pseudo-time using Monocle3 (***Trapnell et al., 2014***). The differentially expressed genes for each trajectory were identified using spatial autocorrelation analysis (***Cao et al., 2019***). The gene ontology terms enriched in each gene module along the trajectories were retrieved in the Gene Ontology Resource database (***Ashburner et al., 2000***; ***Consortium, 2019***) (***Figure 4–Figure Supplement 1***). We estimated the genetic diversity of progenitors for each trajectory using the Watterson estimator, which is given by 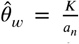, where ***K*** is the number of scarring paths in the trajectory and 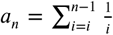 where *n* is the number of cells marked by filtered scarring paths.

### Data availability

Single-cell gene expression data are available in the Sequence Read Archive of the National Center for Biotechnology Information (BioProject ID: PRJNA623798).

## Acknowledgments

We thank J. Huang and Z. Zhang for technical support.

## Additional information

### Author contributions

Zhuoxin Chen, Conceptualization, Methodology, Software, Data curation, Formal analysis, Investigation, Writing – original draft, Writing – review and editing; Chang Ye, Software, Investigation, Resources, Writing – review and editing; Zhan Liu, Investigation, Resources; Shanjun Deng, Software, Resources; Xionglei He, Conceptualization, Writing – review and editing, Supervision, Funding Acquisition; Jin Xu, Conceptualization, Methodology, Writing – original draft, Writing – review and editing, Supervision, Funding Acquisition, Project administration

---

**Figure 1–Figure supplement 1.**
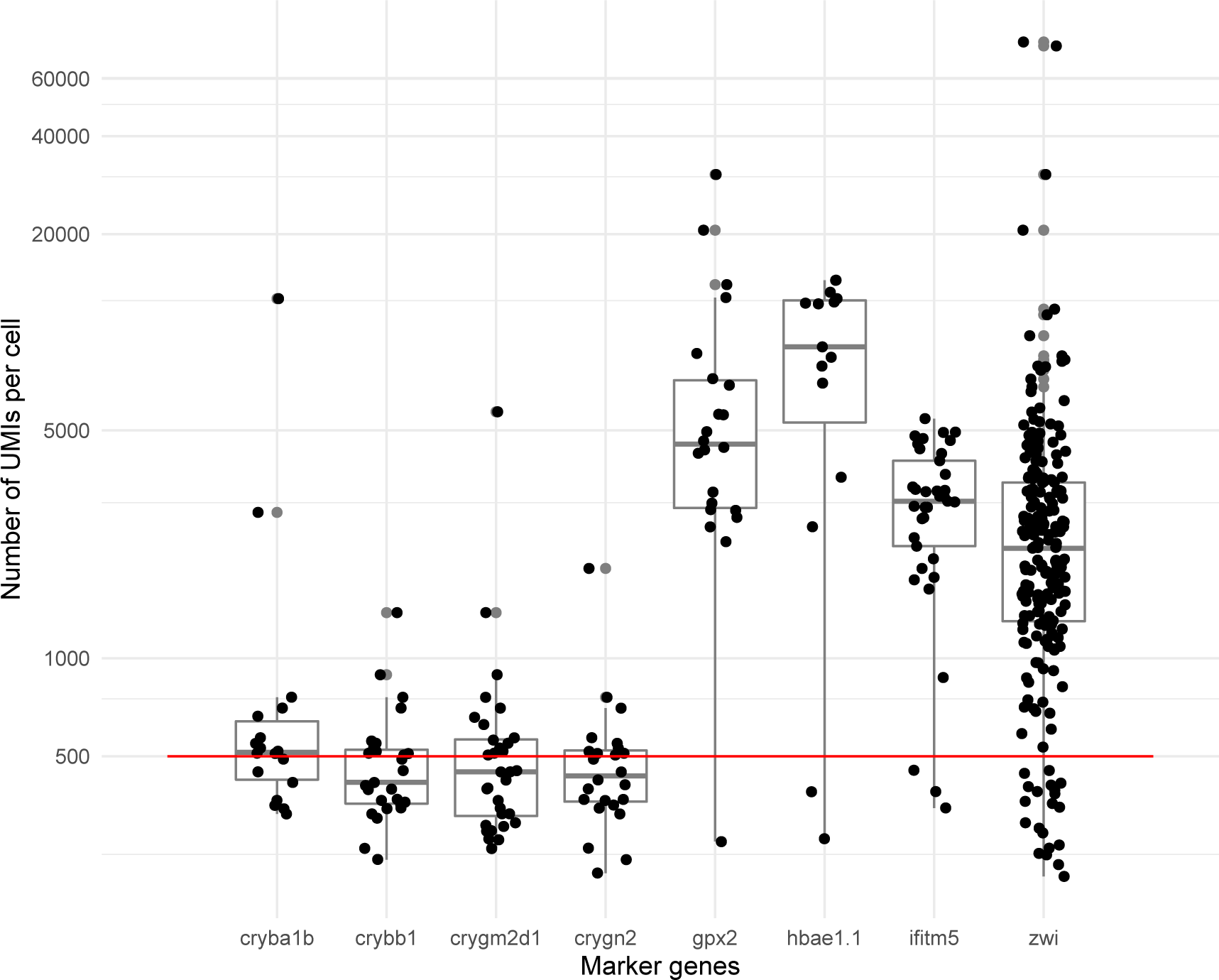
The distribution of UMI number per cell with different marker genes. Most of the cells with lens marker genes (⩾2 UMIs, *cryba1b, crybb1, crygm2d1, crygn2*) had unique reads that were fewer than 500, which is the threshold for consideration as a real cell; whereas a large proportion of cells with other marker genes (⩾2 UMIs, *gpx2, hbae1.1, ifitm5, zwi*) could pass this threshold. Part of the cells with different lens marker genes overlapped.

**Figure 2–Figure supplement 1.**
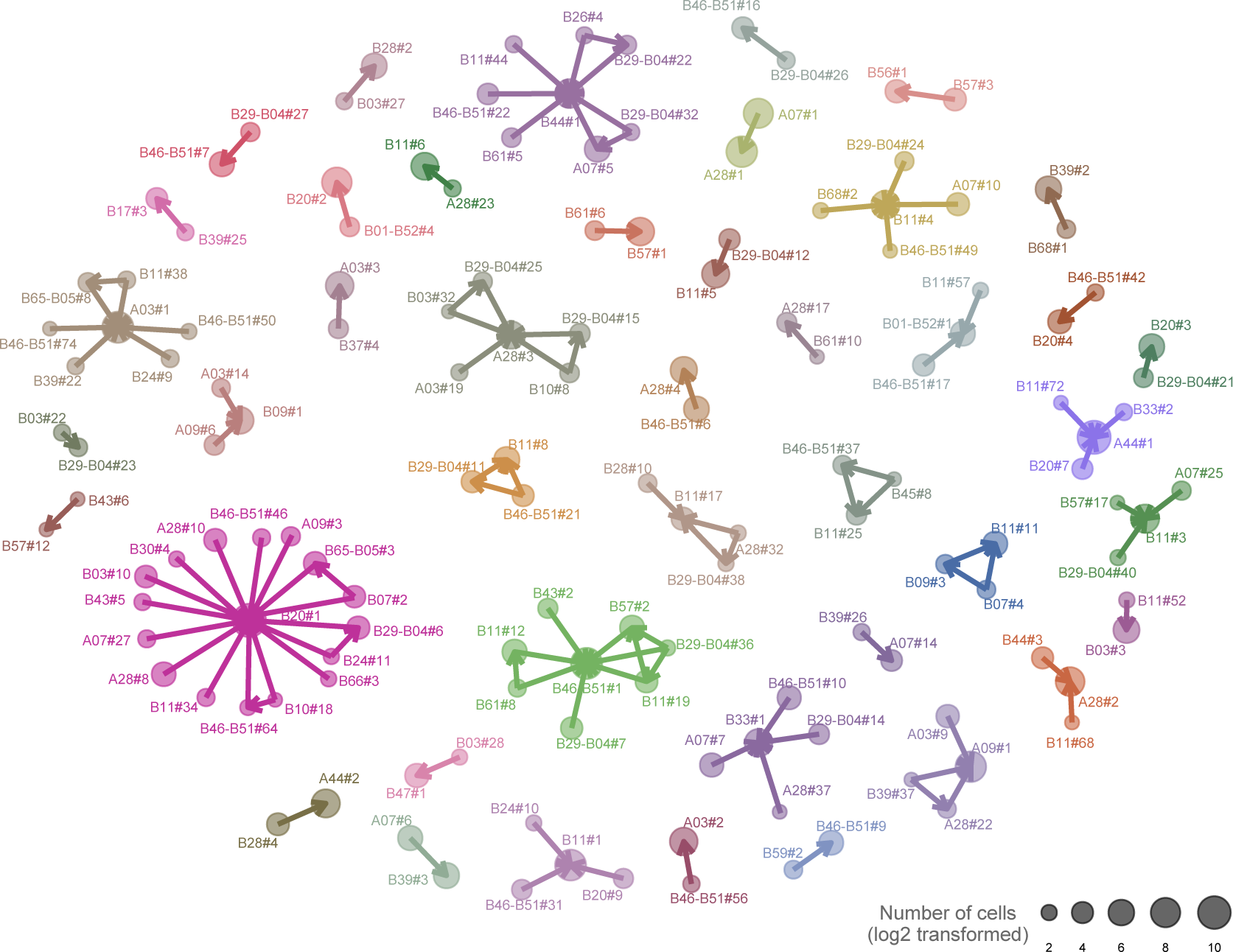
Network relationships of non-recurring scars. The arrows indicate a scar that belongs to the descent to which it points. The size of each point represents the number of cells marked by that scar (*log*_2_ scale). The colors indicate different sub-networks of scar relationships, totaling 42 sub-networks.

**Figure 2–Figure supplement 2.**
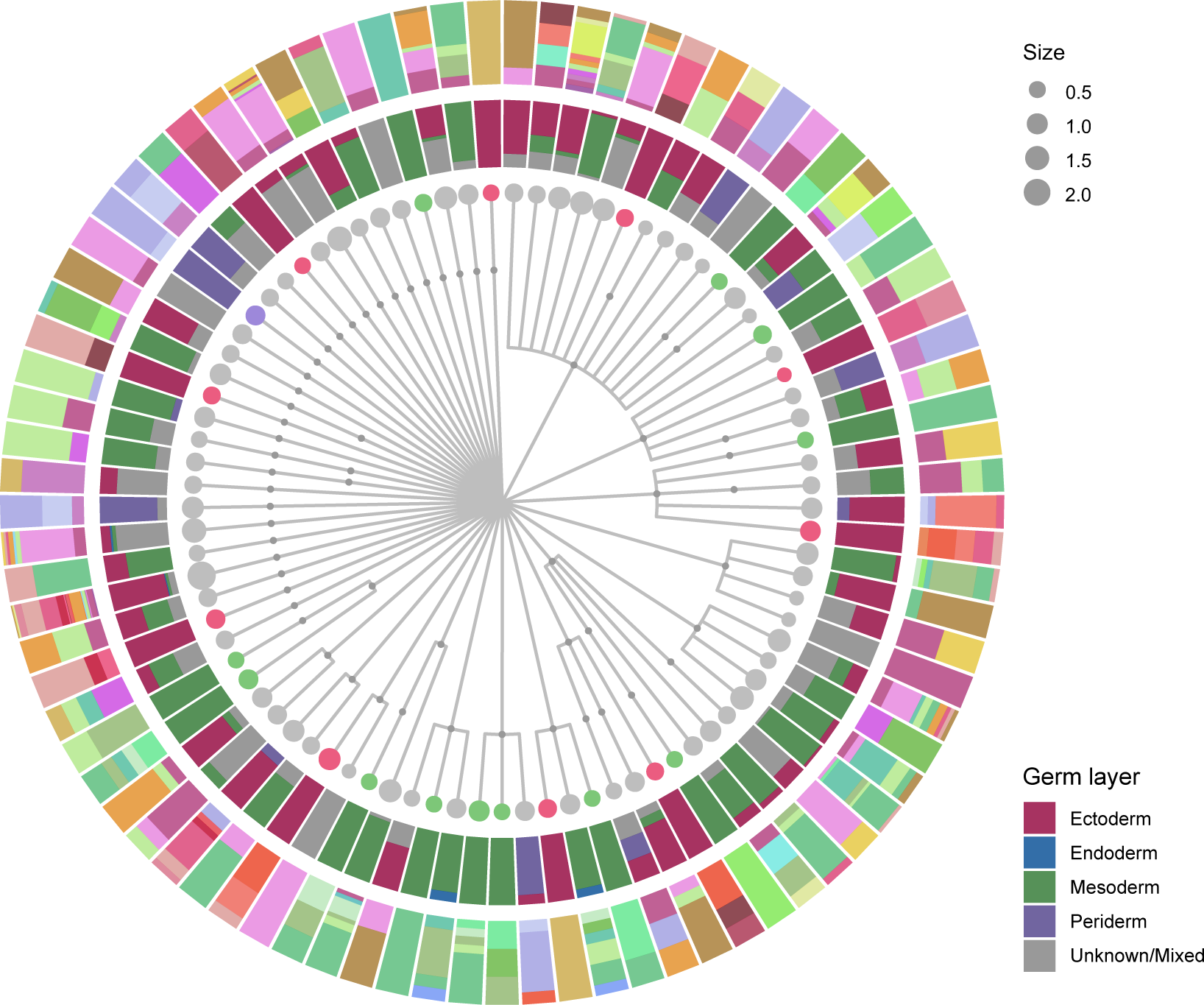
The reconstructed lineage tree for 7 days post-fertilization zebrafish larva. Only unambiguously placed scars were used for lineage reconstruction. Cells were placed to the terminal nodes according to their scar profiles. The terminal node size was scaled by the number of cells within that node (*log*_1_0-transformed) and colored as if it was purely comprised of cells from a single germ layer. The intermediate ring of the histogram refers to the relative proportions of different germ layers. The color code is indicated in the legend. The outer ring of the histogram shows the fractions of cell types within the terminal nodes and the color code is as in Figure 1.

**Figure 2–Figure supplement 3.**
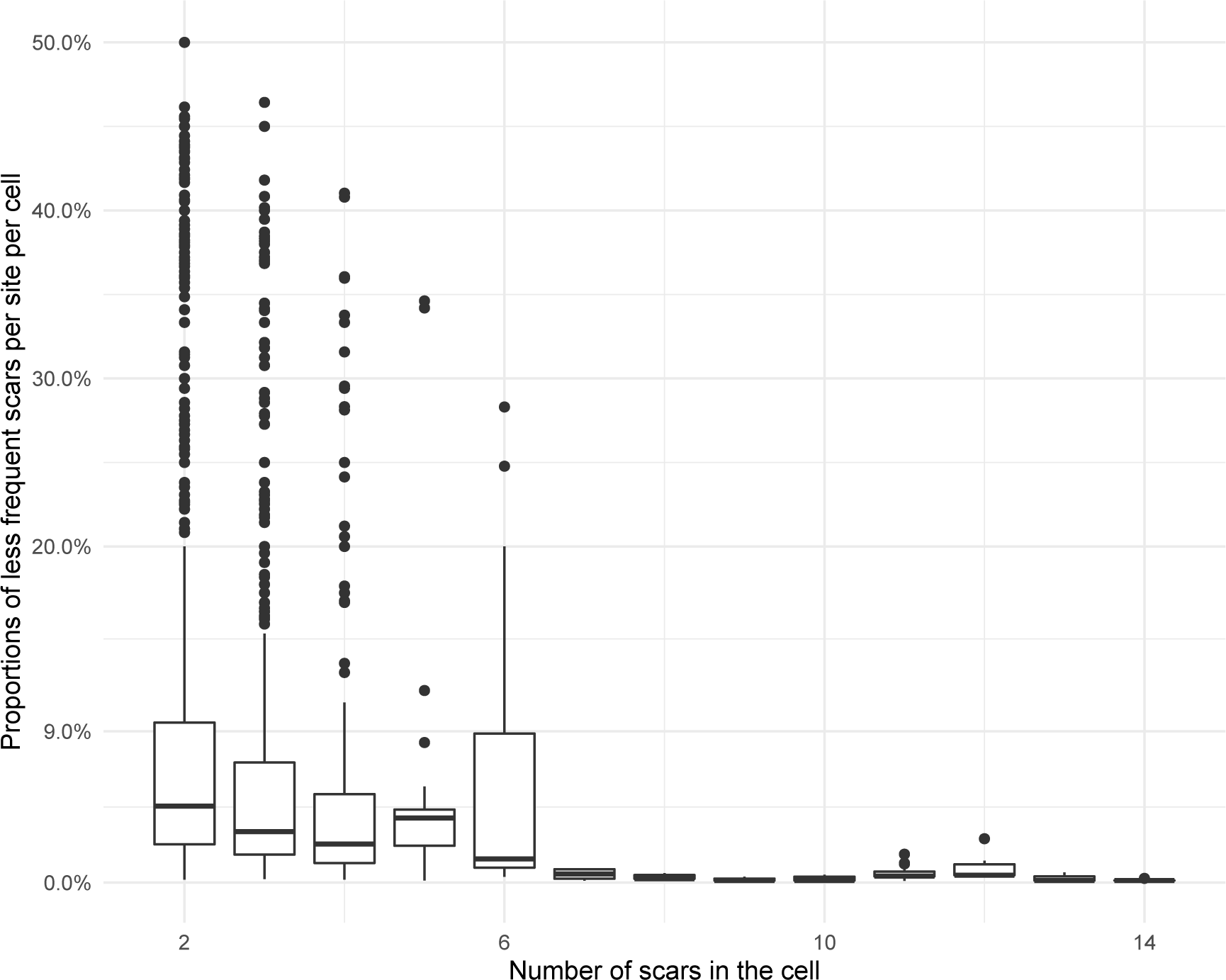
Proportions of less frequent scars per site in individual cells with various number of scars. The 75% quantile of all less frequent scars is ∼9%.

**Figure 2–Figure supplement 4.**
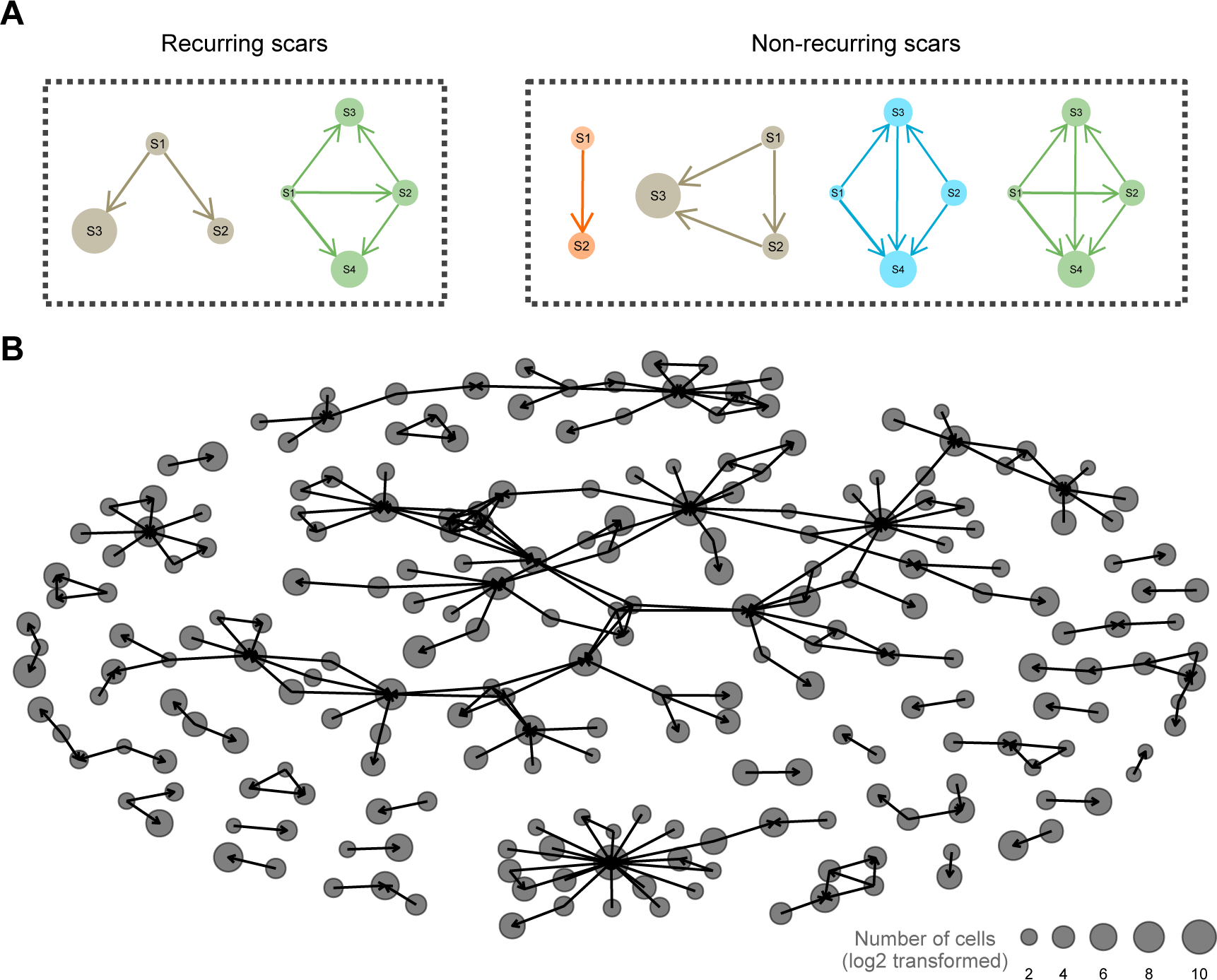
Scar networks before filtering. (**A**) Discrimination of recurring scars and non-recurring scars. (**B**) Scar networks before filtering out recurring scars.

**Figure 3–Figure supplement 1.**
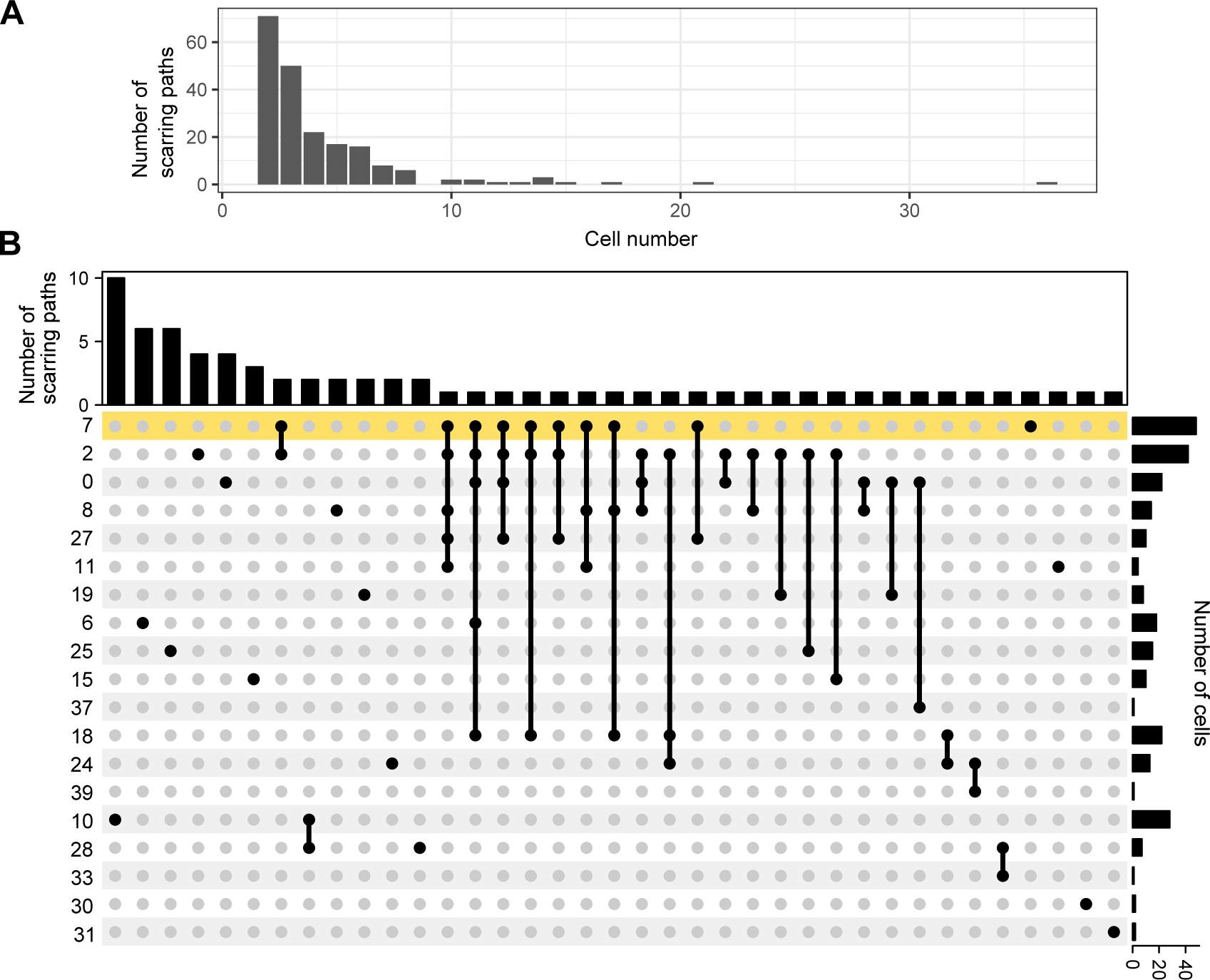
Lineage information for simulation. (**A**) Distribution of the number of cells within each filtered scarring path. (**B**) Upset plot for the number of scarring paths detected within or between cell types from the ectoderm. The yellow shade represents multipotent stem cells.

**Figure 3–Figure supplement 2.**
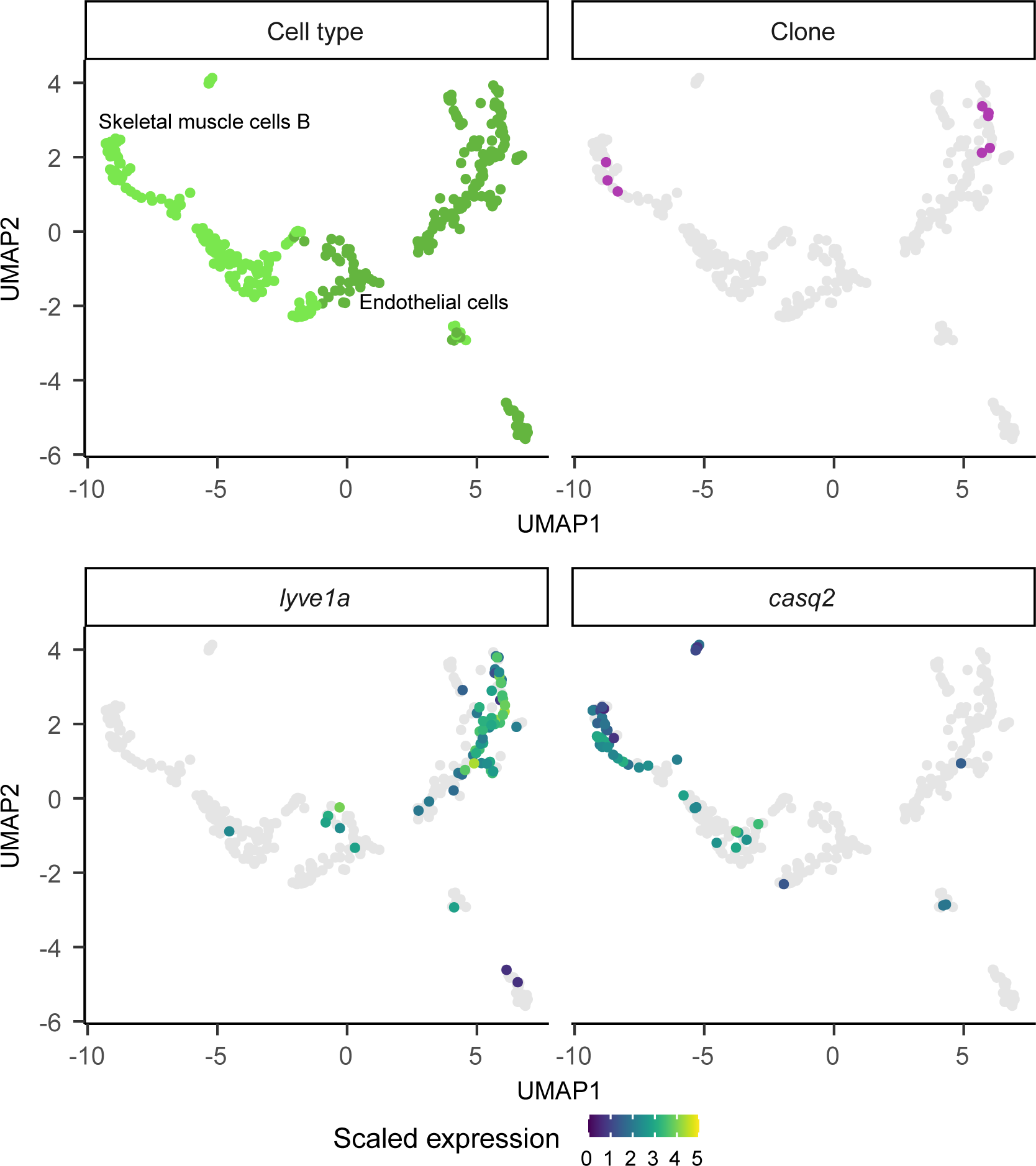
Fin muscle cells share a close relationship with endothelial cells. UMAP visualization of skeletal muscle cells B and endothelial cells (top left) and the cells marked by the same scarring path (top right). The color code for the cell types is as in ***Figure 1***. Expression patterns of marker genes for cardinal vein endothelial cells (lyve1a, bottom left) and fin muscle cells (casq2, bottom right).

**Figure 3–Figure supplement 3.**
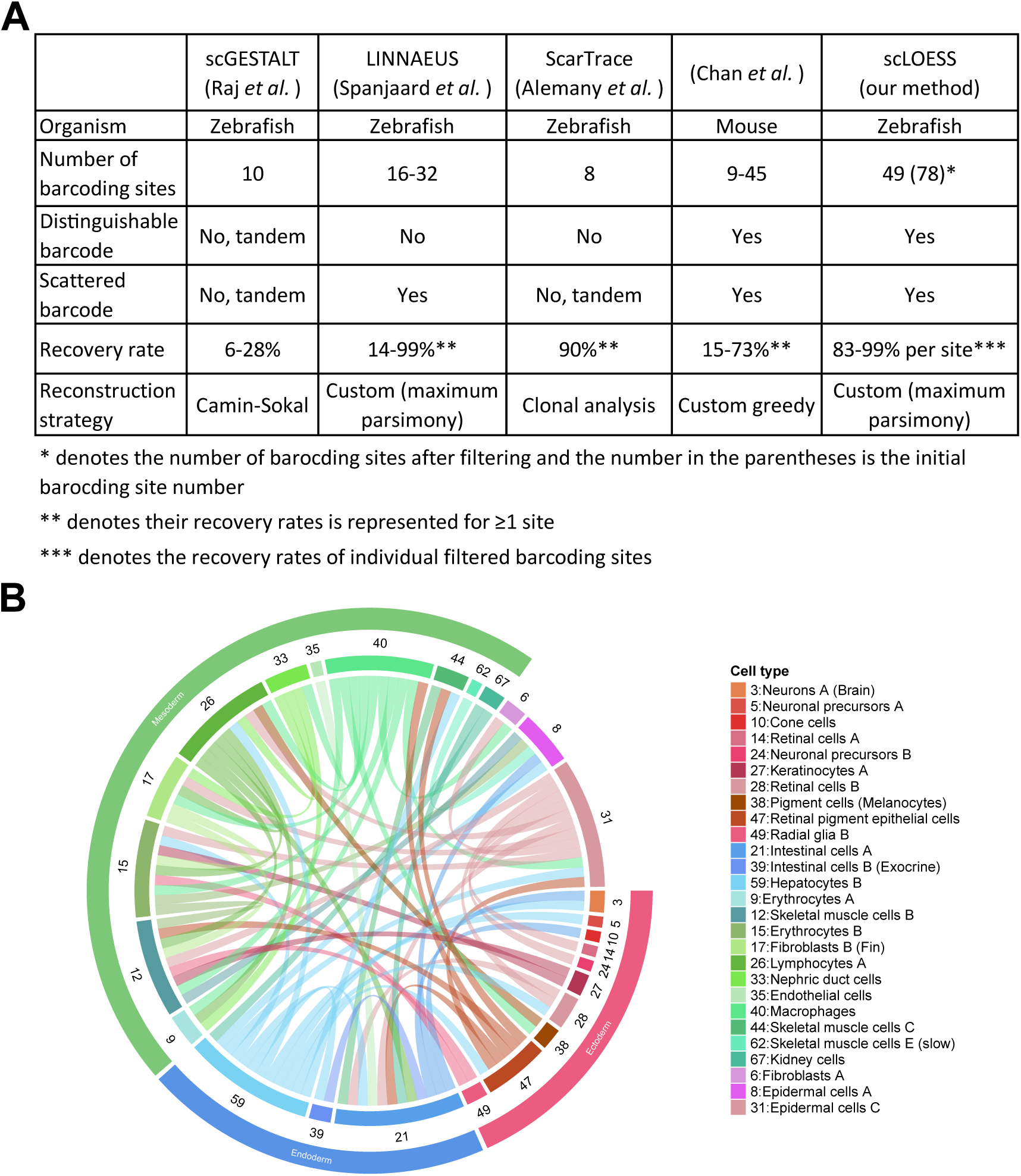
Comparison of various methods for single-cell lineage tracing in vertebrates. (**A**) Table summarizing contemporary Cas9-based lineage recording strategies for vertebrates. (**B**) Chord diagram of cell-type relationships for 5 days post-fertilization larvae (*n* = 7) based on the enrichment of scar connections (only *p*_*adj*_ < 0.01 are shown) between cell types from ***Spanjaard et al.*** (***2018***). This is the same as *Supplementary Figure 8c* in that study, but with a different representation.

**Figure 3–Figure supplement 4.**
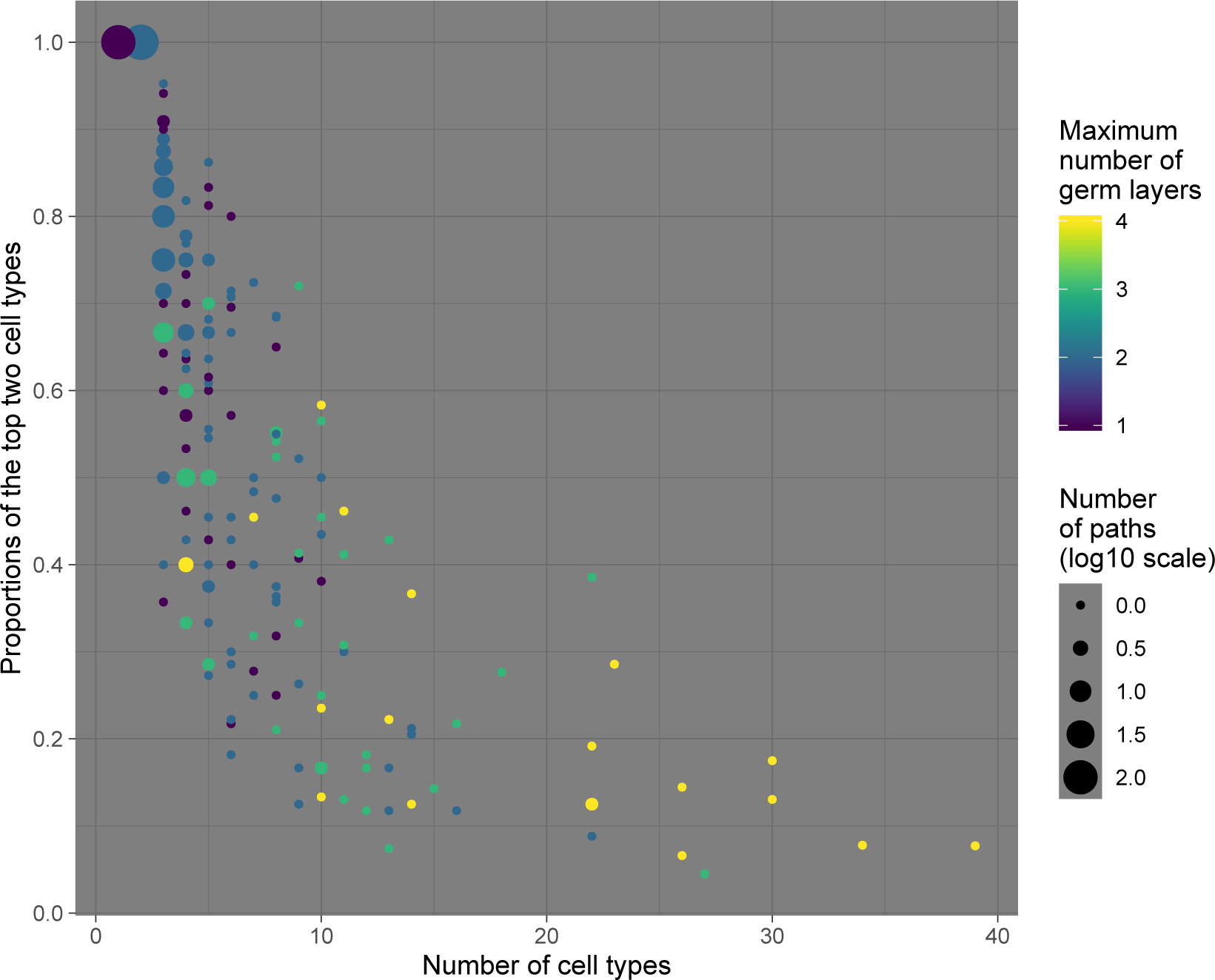
The relationship of the number of cell types and the proportions of the top two cell types in each scarring path. Because different scarring paths may have the same number of cell types and proportions, each point may represent more than one path and the point size is scaled by the number of paths (log10 scale). Points are colored by the maximum number of germ layers of each point.

**Figure 4–Figure supplement 1.**
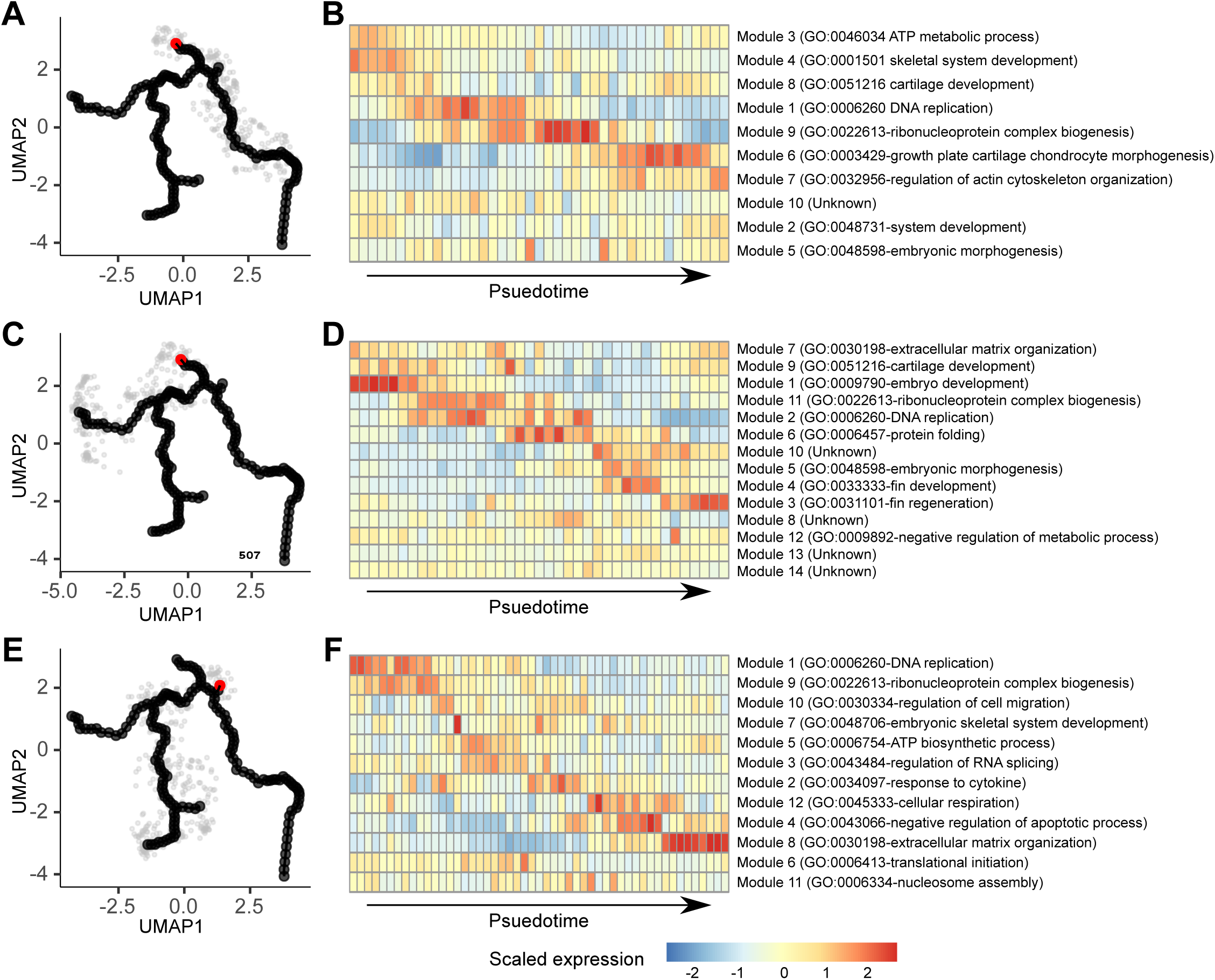
Transcriptional transition along different trajectories. (**A, C, E**) Selected cells along one of the trajectories. The red nodes represent the starting points of each pseudo-time analyses. (**B, D, F**) The gene modules change along the trajectories and the representative gene ontology for each module.

**Figure 4–Figure supplement 2.**
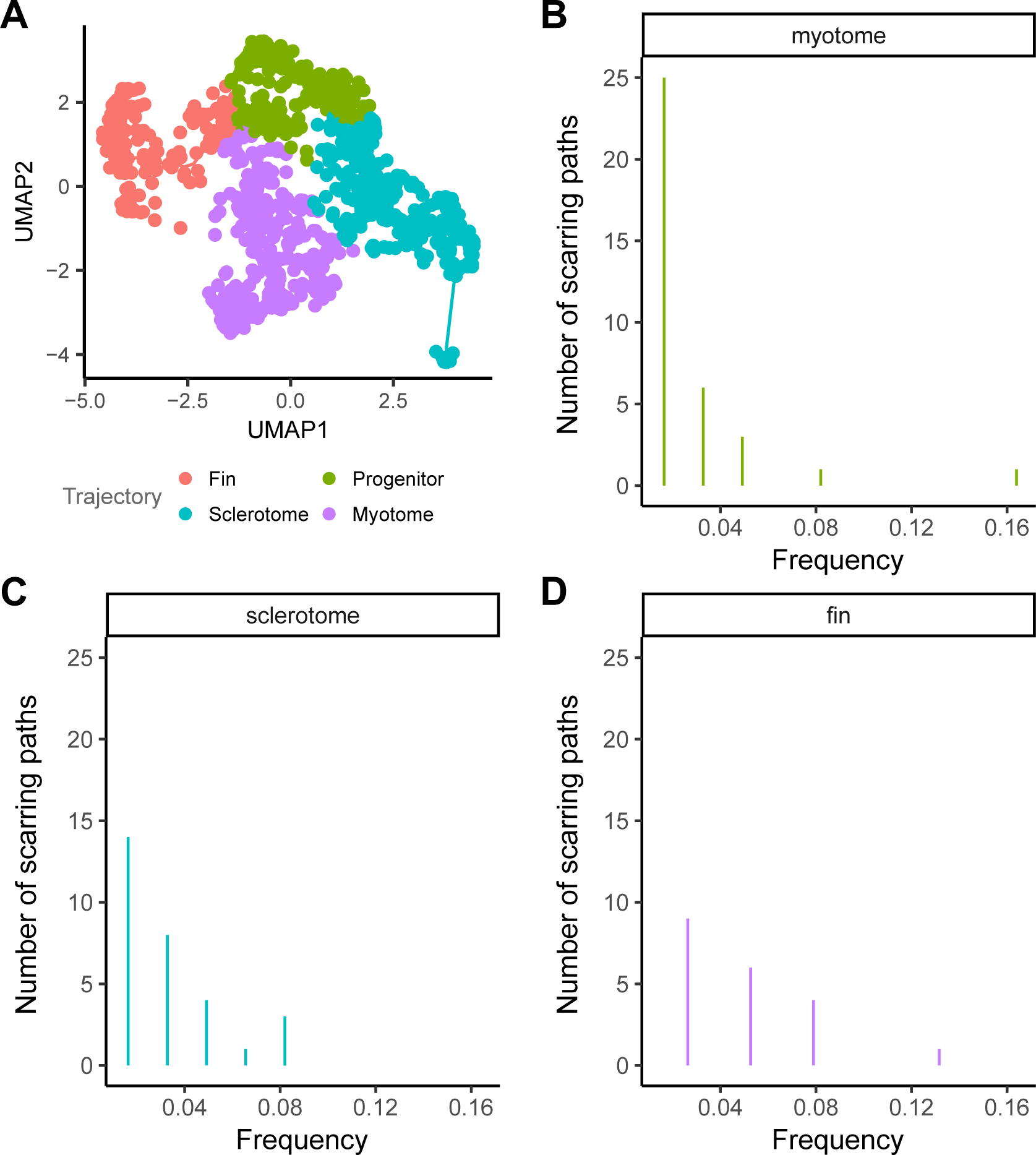
Site frequency spectrum of different trajectories. (**A**) UMAP visualization of cells in the paraxial mesoderm colored by the trajectory to which they belong. (**B**-**D**) Frequency distribution of different scarring paths within distinct trajectories.

